# A novel ENU-induced ankyrin-1 mutation impairs parasite invasion and increases erythrocyte clearance during malaria infection in mice

**DOI:** 10.1101/072587

**Authors:** Hong Ming Huang, Denis C. Bauer, Patrick M. Lelliott, Andreas Greth, Brendan J. McMorran, Simon J. Foote, Gaetan Burgio

## Abstract

Genetic defects in various red blood cell (RBC) cytoskeletal proteins have been long associated with changes in susceptibility towards malaria infection. In particular, while ankyrin (Ank-1) mutations account for approximately 50% of hereditary spherocytosis (HS) cases, an association with malaria is not well-established, and conflicting evidence has been reported. We describe a novel N-ethyl-N-nitrosourea (ENU)-induced ankyrin mutation MRI61689 that gives rise to two different ankyrin transcripts: one with an introduced splice acceptor site resulting a frameshift, the other with a skipped exon. *Ank-1*^*(MRI61689/+)*^ mice exhibit an HS-like phenotype including reduction in mean corpuscular volume (MCV), increased osmotic fragility and reduced RBC deformability. They were also found to be resistant to rodent malaria *Plasmodium chabaudi* infection. Parasites in *Ank-1*^*(MRI61689/+)*^ erythrocytes grew normally, but red cells showed resistance to merozoite invasion. Uninfected *Ank-1*^*(MRI61689/+)*^ erythrocytes were also more likely to be cleared from circulation during infection; the “bystander effect”. This increased clearance is a novel resistance mechanism which was not observed in previous ankyrin mouse models. We propose that this bystander effect is due to reduced deformability of *Ank-1*^*(MRI61689/+)*^ erythrocytes. This paper highlights the complex roles ankyrin plays in mediating malaria resistance.

## Introduction

Malaria is a mosquito-borne disease caused by the protozoan *Plasmodium*, responsible for many deaths every year, mostly children ^1^. In endemic regions with limited healthcare access, host genetics is one of the major determinants of malaria susceptibility and survival ^2–4^. This is evident from the distributions of various genetic polymorphisms in humans, such as Duffy antigen negativity and sickle cell trait, which coincide with malaria distribution ^3,5,6^. It is thought that these genetic polymorphisms confer protection against malaria, thus providing a survival advantage in the face of malaria-induced mortality ^7,8^.

In addition, these polymorphisms also provide crucial insights into host-parasite interactions. *Plasmodium* relies on a favourable host environment in order to thrive, many erythrocyte-related polymorphisms have been discovered that interfere with parasite survival thus contributing to malaria resistance. These include polymorphisms that affect the cytoskeleton of erythrocytes, such as Southeast Asian Ovalocytosis (SAO), hereditary elliptocytosis (HE) and spherocytosis (HS) ^9–13^. Several hypotheses have been proposed for the mechanisms by which they confer malaria protection, including reduced erythrocyte invasion, intra-erythrocytic growth and cytoadherence ^8,13–19^. However, due to the heterogeneity of the manifestation of these disorders in the human population, contradicting evidences for resistance mechanisms has often been presented. A study done by Facer ^13^ showed that only patients carrying certain spectrin mutations have impaired parasite invasion of red blood cells (RBC), but not others that also exhibited HE symptoms. A similar observation was reported by Chishti, et al. ^17^, where individuals with defective protein 4.1 exhibited intra-erythrocytic growth inhibition, but not those with glycophorin C defects, despite the fact that both defects gave rise to HE. These differences in malaria resistance mechanisms remained largely unexplored, and further studies in this aspect would potentially provide useful insight into host-parasite interactions.

The RBC cytoskeletal protein ankyrin-1 (ANK-1), is a 210kDa protein responsible for connecting the spectrin network with the RBC membrane through interactions with Band 3, protein 4.2 and the Rhesus complex ^20–22^. Spherocytosis is a genetic disorder where RBCs are abnormally small and are known as spherocytes. ANK-1 mutations account for more than 50% of human HS cases ^23^. However, similar to SAO and HE, HS is a heterogeneous disorder where the symptoms vary greatly depending on the mutations. The disorder can range from asymptomatic through to severe anaemia requiring splenectomy ^24^. Despite a possible association with malaria, HS is actually common in Northern European and Japanese populations with frequency of about 1 in 2000 individuals ^25–28^, but much rarer in other populations ^29^. Nevertheless, several *in vitro* and *in vivo* studies have repeatedly reported association between HS and malaria resistance, and several mechanisms have been suggested. An *in vitro* study done by Schulman, et al. ^16^ using RBCs from HS patients suggested that parasite invasion and growth in these erythrocytes was impaired. This is further supported by studies done in mice with ankyrin mutations. Both *Ank-1*^*(nb/nb)*^ and *Ank-1* ^*(MRI23420/+)*^ have shown inhibited intra-erythrocytic growth and erythrocyte invasion possibly due to spectrin and ankyrin deficiency, respectively ^30,31^. On the other hand, another mutation described by Rank, et al. ^32^, *Ank-1*^*1674/+*^, parasite invasion appeared to be normal in these erythrocytes. Instead, increased erythrocyte fragility was proposed as a contributing factor for increased malaria resistance ^32^. Taking these observations together, it is possible that disruption to erythrocyte cytoskeletons can mediate multiple mechanisms of resistance.

In a large phenotypic N-ethyl-N-nitrosourea (ENU) mutagenesis screen, using either abnormal red cells or resistance to malaria as to the screened phenotypes, we identified many novel mutations that give rise to RBC abnormalities and consequently malaria resistance in mice. We report here an ENU-induced mutation in the ankyrin-1 gene (*Ank-1*^*(MRI61689)*^) which was found to exhibit a HS-like phenotype, with significantly lower RBC volumes, increased osmotic fragility and decreased deformability. *Ank-1*^*(MRI61689)*^ also confers resistance towards *Plasmodium chabaudi adami* infection in mice, and *Ank-1*^*(MRI61689/+)*^ mice were shown to show both reduced merozoite invasion and increased RBC clearance, possibly as a consequence of reduced red blood cell deformability.

## Results

### The MRI61689 mutation gives rise to an hereditary spherocytosis-like phenotype

The G1 mouse carrying the MRI61689 mutation was initially identified from an ENU suppressor screen for the recessive mutation db/db. The G1 MRI61689 exhibited abnormal blood parameters on an ADVIA haematological analyser, with reduced MCV of 48.6fL compared to the background of 53.3±0.5fL. This MCV values from the B6.BKS(D)-Lepr^db^/J background is comparable to C57BL/6 mice. We crossed the G1 founder mouse with B6.BKS(D)-Lepr^db^/J to produce G2 mice where approximately half of the animals exhibited an abnormal phenotype (Table 1). The affected G2 progeny, which were obligate heterozygotes for the ENU-induced mutation, showed reduction in MCV (46.1±0.2fL) compared to unaffected progeny (51.4±0.4fL), lower mean corpuscular haemoglobin (MCH) (13.5±0.1pg compared to 14.6±0.1pg of wild-type), elevated RBC count (11.1±0.1x10^9^ cells/ml compared to 10.5±0.1x10^9^ cells/ml of wild-type) (Table 1). No differences were observed for total haemoglobin (HB), mean corpuscular haemoglobin concentration (MCHC), white blood cell (WBC) count, platelets count (PLT) or reticulocyte percentage (Table 1).

**Table 1.** The complete blood count of *Ank-1*^*(MRI61689/+)*^ mice. The haematological parameters of *Ank-1*^*(MRI61689/+)*^ compared to wild-type mice (n=19-25). WBC = white blood cell count; RBC = red blood cell count; HGB = haemoglobin; MCV = mean corpuscular volume; MCH = mean corpuscular haemoglobin; MCHC = mean corpuscular haemoglobin concentration; PLT = platelet concentration; -Retics = percentage of reticulocytes.

|  | WBC<br>(x10<sup>6</sup> /ml) | RBC<br>(x10<sup>9</sup> /ml) | HGB<br>(g/L) | MCV<br>(fL) | MCH<br>(pg) | MCHC<br>(g/L) | PLT<br>(x10<sup>6</sup> /ml) | %Retics |
| --- | --- | --- | --- | --- | --- | --- | --- | --- |
| Wild type | 8.7±0.5 | 10.5±0.1 | 153.2±2.2 | 51.4±0.4 | 14.6±0.1 | 283.6±2.8 | 1151±57 | 2.69±0.26 |
| *Ank-1*<sup>(MRI61689/+)</sup> | 11.9±4.5 | 11.1±0.1 | 151.2±1.4 | 46.1±0.2 | 13.5±0.1 | 281.4±9.3 | 1151±61 | 2.24±0.21 |
| p-values | NS | P<0.01 | NS | P<0.001 | P<0.001 | NS | NS | NS |

However, when two affected mice were intercrossed, a quarter of the pups were found to die within 1 week postnatally, suggesting homozygosity for MRI61689 might be incompatible with life. Blood smears were taken from these pups and compared with the other affected and unaffected mice. The heterozygotes have slightly smaller RBCs but no target cells or spherocytes were observed (Figure 1a). Conversely, homozygous mice had significantly smaller RBCs with anisocytosis, fragmented RBCs, acanthocytes and reticulocytosis (Figure 1a). Under SEM, RBCs of heterozygous mice seemed to have less distinct discoid shape, but otherwise no distinguishing features were observed. (Figure 1b). On the other hand, the RBCs of homozygous mice appeared very deformed, acanthocytic and appeared to lack the discoid shape (Figure 1b).

**Figure 1.**
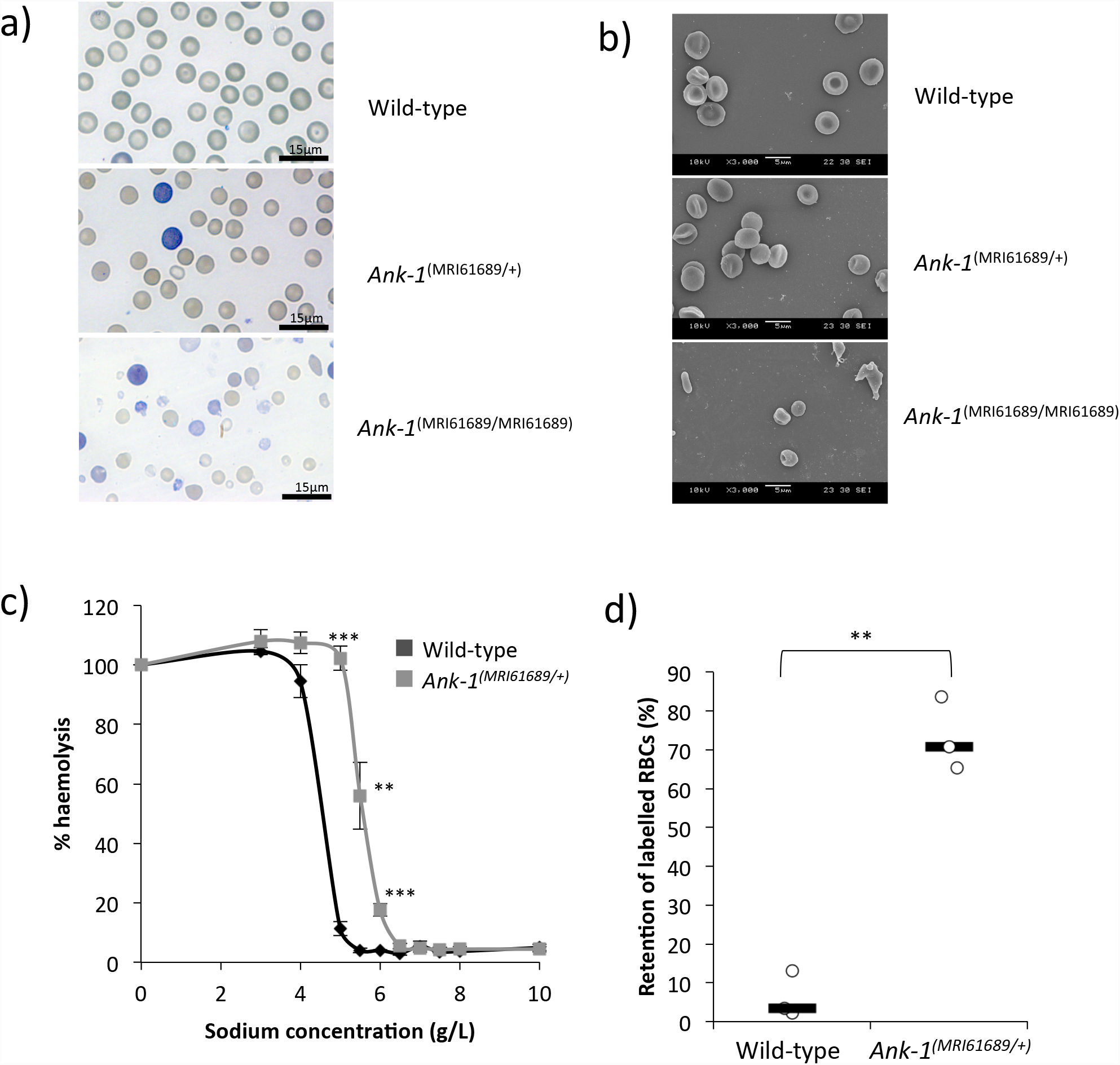
The phenotypic characterisation of *Ank-1*^*(MRI61689/+)*^ mice. The morphology of *Ank-1*^*(MRI61689/+)*^ and *Ank-1*^*(MRI61689/MRI61689)*^ erythrocytes under light microscopy with Giemsa stain (a) and scanning electron microscopy (b). The osmotic fragility curve of wild-type and *Ank-1*^*(MRI61689/+)*^ erythrocytes when subjected to osmotic stress (c) (n=5-7 per group). The *in-vitro* spleen retention rate of wild-type and *Ank-1*^*(MRI61689/+)*^ erythrocytes when passing through filter beds (d) (n=3 per group). ** indicates P<0.01, *** indicates P<0.001, and all error bars are standard error of mean (SEM).

When subjected to osmotic stress, RBCs of heterozygous mice showed significantly increased fragility compared to wild-type erythrocytes (with 50% haemolysis at approximately 5.6 g/L compared to 4.5 g/L of wild-type) (Figure 1c). The RBC deformability was assessed using an *in vitro* spleen retention assay by filtering RBCs through a layer of beads with varying sizes. This is thought to model splenic filtration *in vivo*, with retention thought to indicate reduced deformability. As shown in Figure 1d, up to 70% of the RBCs from heterozygous MRI61689 mice were retained within the bead layer compared to 3.5% of wild-type RBCs, suggesting a significantly reduced RBC deformability in the presence of this ankyrin mutation.

### MRI61689 carries a splice site mutation in *Ank-1* gene resulting in an alternative transcript and exon skipping

To identify the causative mutation responsible for this abnormal RBC count, we sequenced the exomes of 2 heterozygous mice. Exome sequencing revealed a number of variants. These were prioritised based on filters as shown in Table 2. Through further genotyping using Sanger sequencing, a mutation in *Ank-1* gene was found to correlate with all the affected mice and was proven to segregate perfectly with the reduced MCV for over 3 generations of mouse crosses. The mutation was found in the 17-18 intron of *Ank-1* gene, with T to A transversion 11 base pair upstream of exon 18 (IVS17-11T>A) (Figure 2a). This is situated in the ankyrin-repeats domain involved in band 3 binding. We proposed that the mutation introduced a new acceptor splice site for exon 18, potentially leading to a frameshift mutation.

**Figure 2.**
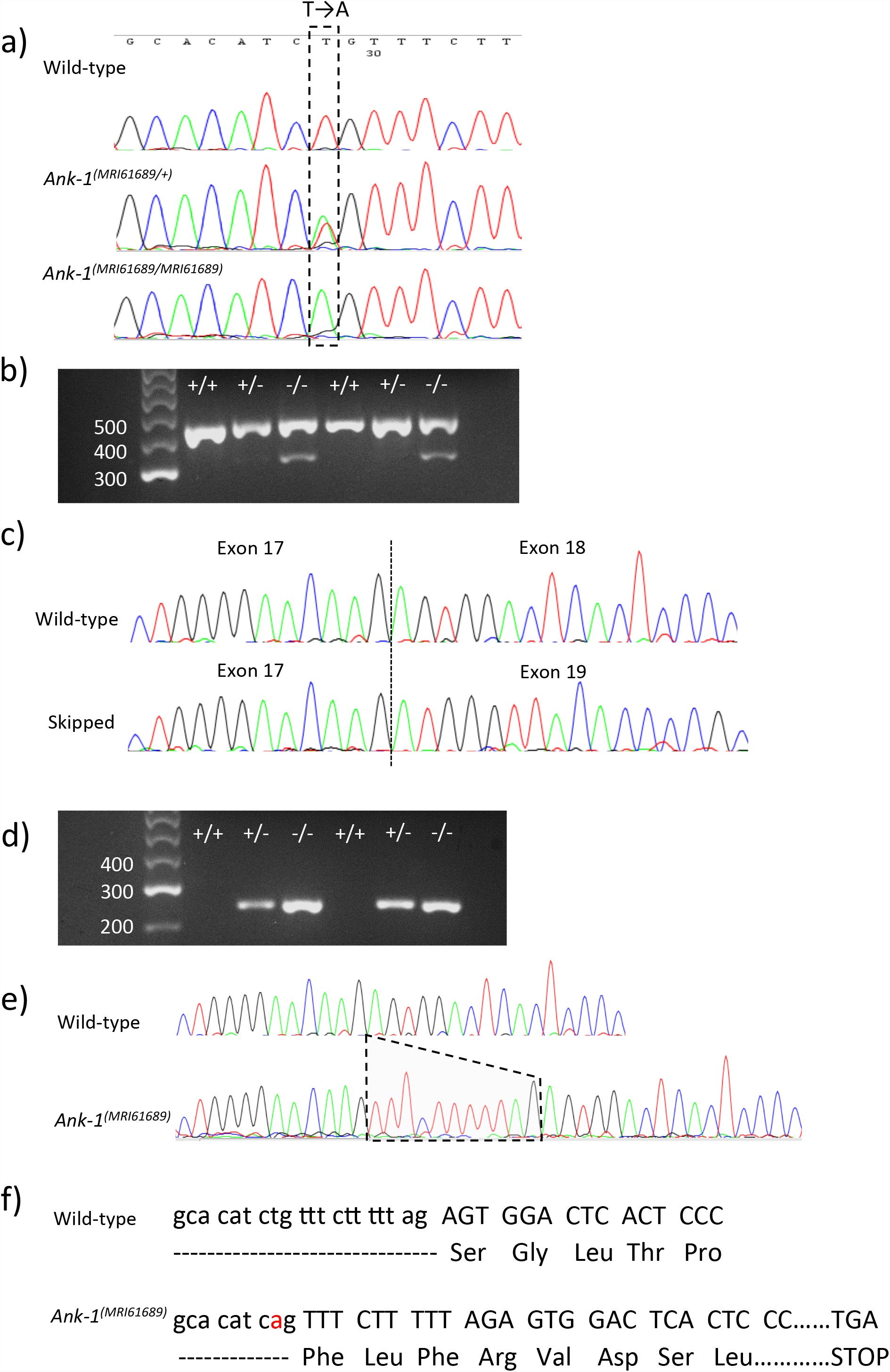
The identification of *Ank-1*^*(MRI61689)*^ mutation and its effects on transcription. The sequencing of *Ank-1*^*(MRI61689)*^ mutation, showing a T to A transversion (a). Gel electrophoresis of amplified cDNA product from wild-type, *Ank-1*^*(MRI61689/+)*^ and *Ank-1*^*(MRI61689/MRI61689)*^ embryonic livers with primers that spanned the exon 17 to 21 of ankyrin-1 cDNA (primer set 1) (b). The sequencing results of both bands, showing exon skipping in the abnormal transcript of *Ank-1*^*(MRI61689/MRI61689)*^ embryonic liver (c). Gel electrophoresis of amplified cDNA product from wild-type, *Ank-1*^*(MRI61689/+)*^ and *Ank-1*^*(MRI61689/MRI61689)*^ embryonic livers with primers that contained the predicted 11bp insertion (primer set 2) (d). Sequencing result showing an 11bp insertion between exon 17 and 18 cDNA of the *Ank-1*^*(MRI61689)*^ transcript (e). The predicted effect of the insertion on the translation of ankyrin-1, showing a frameshift and a premature chain termination (f).

**Table 2:**
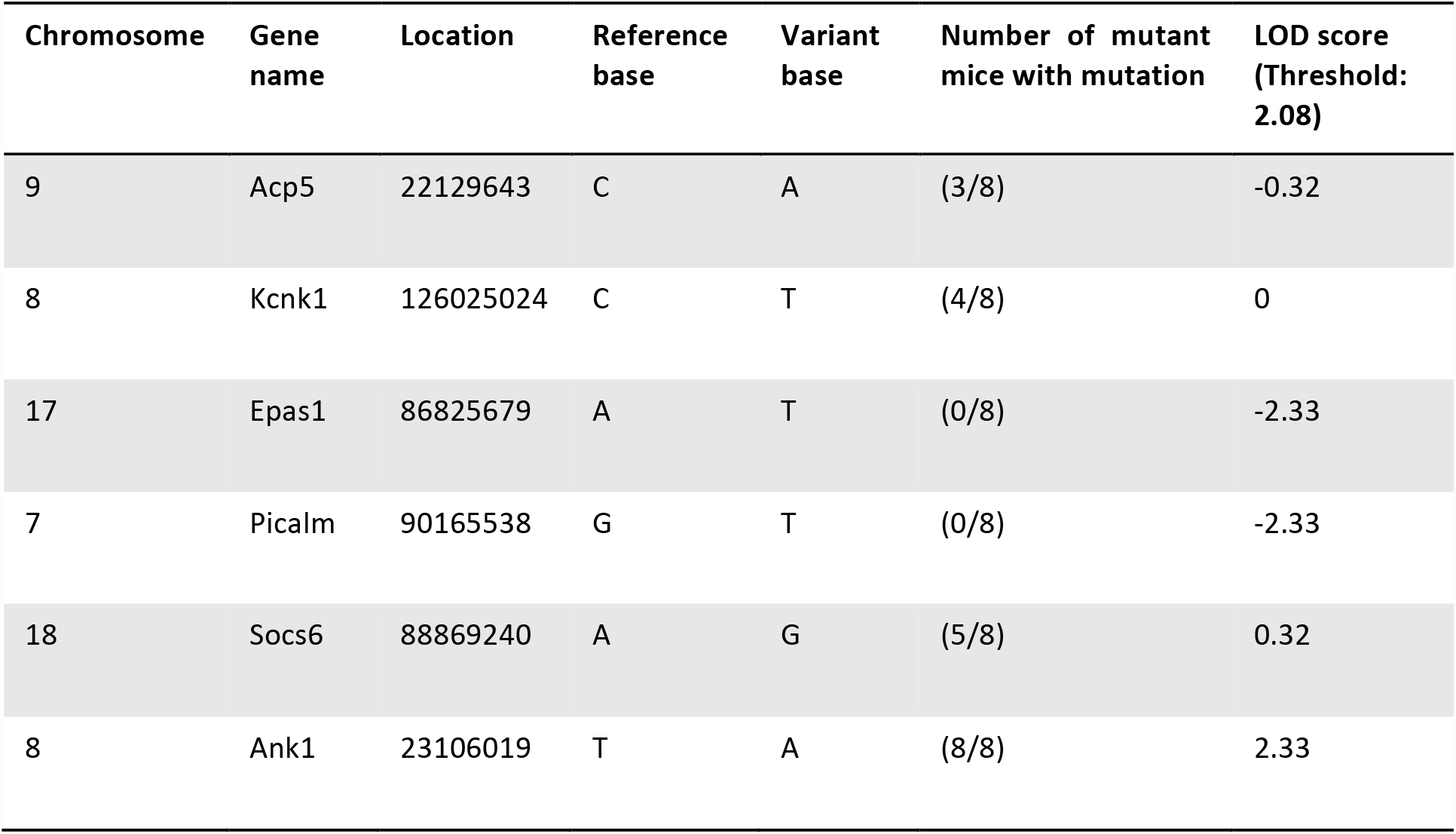
The identification of MRI61689 mutation. Six variants were selected from the exome sequencing, each mutation was sequenced in *Ank-1*^*(MRI61689/+)*^ mice and the number of mutant mice carrying each mutation was determined and LOD score was calculated, with LOD threshold being 2.08 (n=8).

| Chromosome | Gene name | Location | Reference base | Variant base | Number of mutant mice with mutation | LOD score (Threshold: 2.08) |
| --- | --- | --- | --- | --- | --- | --- |
| 9 | Acp5 | 22129643 | C | A | (3/8) | -0.32 |
| 8 | Kcnk1 | 126025024 | C | T | (4/8) | 0 |
| 17 | Epas1 | 86825679 | A | T | (0/8) | -2.33 |
| 7 | Picalm | 90165538 | G | T | (0/8) | -2.33 |
| 18 | Socs6 | 88869240 | A | G | (5/8) | 0.32 |
| 8 | Ank1 | 23106019 | T | A | (8/8) | 2.33 |

To assess this hypothesis, transcript analysis was performed. Embryonic liver RNA was extracted, cDNA was synthesized and PCR-amplified using primers listed in the experimental procedures. Figure 2b shows the PCR products of embryonic liver cDNA from non-mutant, *Ank-1*^*(MRI61689/+)*^ and *Ank-1*^*(MRI61689/MRI61689)*^ when amplified using primer set 1, which covers exon 17 to 21. Bands of approximately 400bp can be observed in all the genotypes, but *Ank-1*^*(MRI61689/MRI61689)*^ also exhibited a second smaller product of approximately 300bp length. We proposed that this second band resulted from exon skipping. Sanger sequencing of these PCR products revealed that the 300bp product lacked exon 18, confirming that the exon 18 was skipped, and exon 19 was directly connected to exon 17 during transcription (Figure 2c). This transcript is predicted to produce a shortened, in-frame 207kDa ANK-1 protein.

To examine the effect of MRI61689 mutation in heterozygous mice, we further designed a primer set containing the predicted acceptor splice site (primer set 2). Figure 2d shows that the mutant transcript is only present in *Ank-1*^*(MRI61689/+)*^ and *Ank-1*^*(MRI61689/MRI61689)*^ mice, as predicted. Further Sanger sequencing revealed an insertion of 11bp into the transcript adding an additional donor splicing site and causing a frameshift mutation in the exon that would result in a premature stop codon at amino acid position 724, as illustrated in Figure 2e, thus giving rise to a truncated protein of 78.5kDa. Therefore the homozygous mice exhibit a mutation at 11 bp upstream of the exon 18 donor splicing site resulting in two alternative transcripts: the skipping of exon 18 and an 11bp insertion and creation of an additional donor splicing site leading to frameshift mutation that would result in a premature stop codon and a truncated protein.

We hypothesize that this mutation would reduce the *Ank-*1 expression levels. We assessed this hypothesis by examining the gene expression levels of *Ank-1* in embryonic liver using qPCR at mRNA level and Western blotting at protein level in mature RBCs. As shown in Figure 3a, *Ank-1* mRNA levels in both *Ank-1*^*(MRI61689/+)*^ and *Ank-1*^*(MRI61689/MRI61689)*^ E14 embryonic livers were significantly reduced, up to 60% and 80% reduction compared to the wild-type, respectively. However, no significant reduction in the full length ANK-1 (210kDa) protein levels was observed in both Coomassie and Western blotting (Figure 3b-d). No truncated form of ANK-1 (78.5kDa) was observed in *Ank-1*^*(MRI61689/+)*^ erythrocytes. Furthermore, no reduction was observed for the protein levels of other cytoskeletal proteins, including Band 3, protein 4.2, alpha- and beta-spectrin (Figure 3b and 3c, Supp. Figure 1). This suggested that erythrocyte protein levels might be compensated by the WT allele in *Ank-1*^(MRI61689/+)^ mice, and the reduction in *Ank-1* mRNA levels did not seem to affect the protein levels.

**Figure 3.**
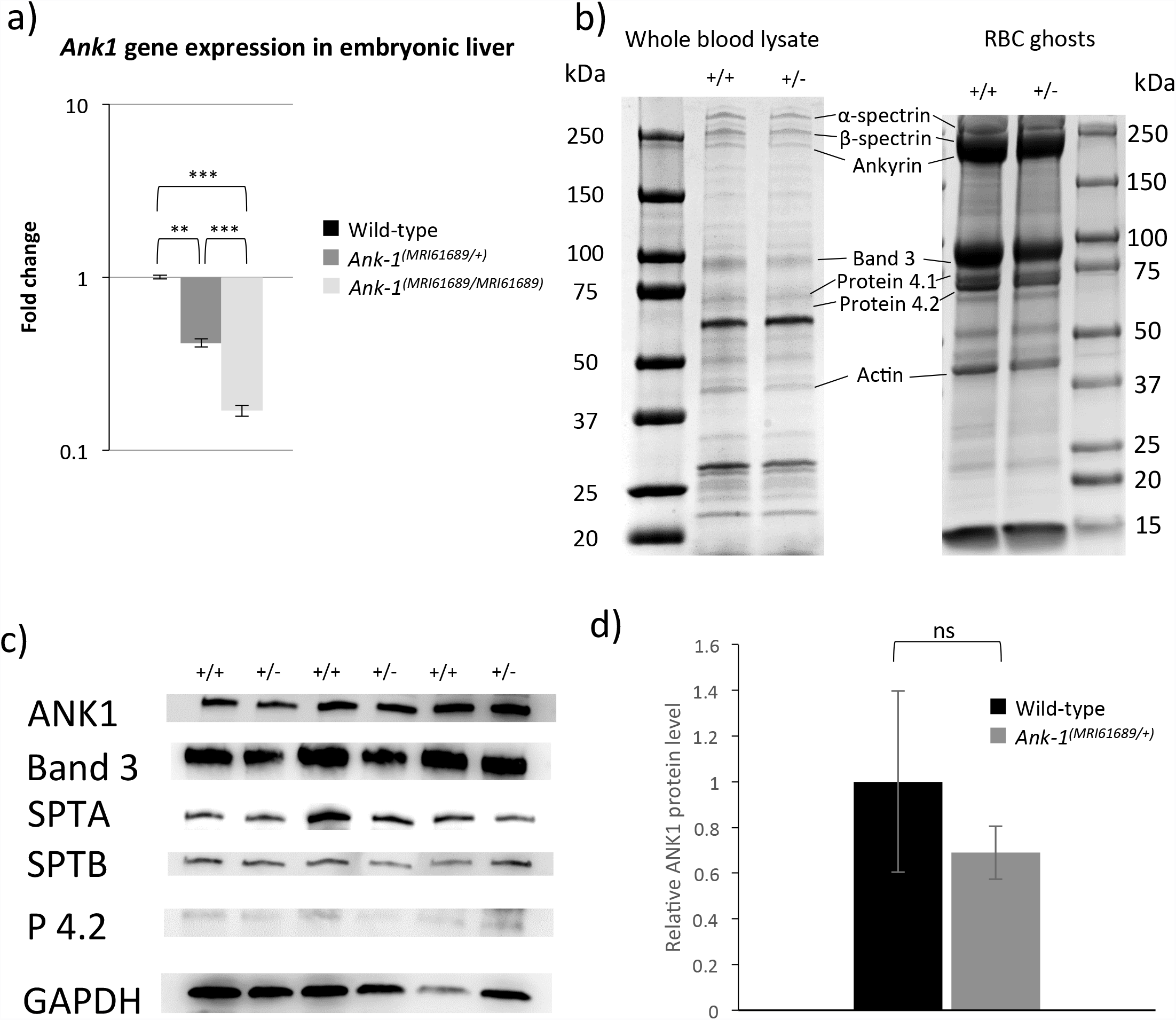
The effect of *Ank-1*^*(MRI61689)*^ mutation on the *Ank-1* expression. Quantitative PCR showing the ankyrin-1 mRNA levels in both *Ank-1*^*(MRI61689/+)*^ and *Ank-1*^*(MRI61689/MRI61689)*^ embryonic liver (a). The protein levels of various cytoskeletal proteins examined with both Coomassie (b) and Western blot (c). The relative protein levels of ankyrin-1 calculated from the western blot (d) (n=3 per group). Error bars indicate SEM.

### *Ank-1*^*(MRI61689/+)*^ mice are resistant to *Plasmodium chabaudi* infection

We hypothesized that the *Ank-1*^*(MRI61689)*^ mutation confers malaria resistance. The malaria susceptibility of *Ank-1*^*(MRI61689/+)*^ mice was examined by injecting a lethal dose of *Plasmodium chabaudi adami DS*, a murine strain of malaria that models the *Plasmodium falciparum* erythrocytic stage ^33^. The *Ank-1*^*(MRI61689/+)*^ mice exhibited significantly lower peak parasitemia, with only approximately 13% parasitemia compared to 52% parasitemia of wild-type (Figure 4a) but no delay in the appearance of parasites was observed. In addition, *Ank-1*^*(MRI61689/+)*^ have a significantly increased survival rate, where all the *Ank-1*^*(MRI61689/+)*^ mice survived the infection (Figure 4b) compared to the 16% survival of wild-type mice. Since *Ank-1*^*(MRI61689)*^ directly affects the red cell (cytoskeletal protein), we hypothesized that the resistance was likely due to a RBC-autonomous effect. Therefore, we postulated three mechanisms of *P. chabaudi* resistance in *Ank-1*^*(MRI61689/+)*^ mice. Firstly, the maturation of parasite inside the *Ank-1*^*(MRI61689/+)*^ erythrocytes could be impaired leading to reduced growth and death of parasites ^31^. Secondly, the *Ank-1*^*(MRI61689/+)*^ erythrocytes might be resistant to merozoite invasion, which resulted in reduced parasitemia and delayed course of infection ^15^. Finally, the infected *Ank-1*^*(MRI61689/+)*^ erythrocytes might more prone to destruction during the course of infection (increased clearance), thus posing a challenge for the parasite to establish a successful infection ^34^.

**Figure 4.**
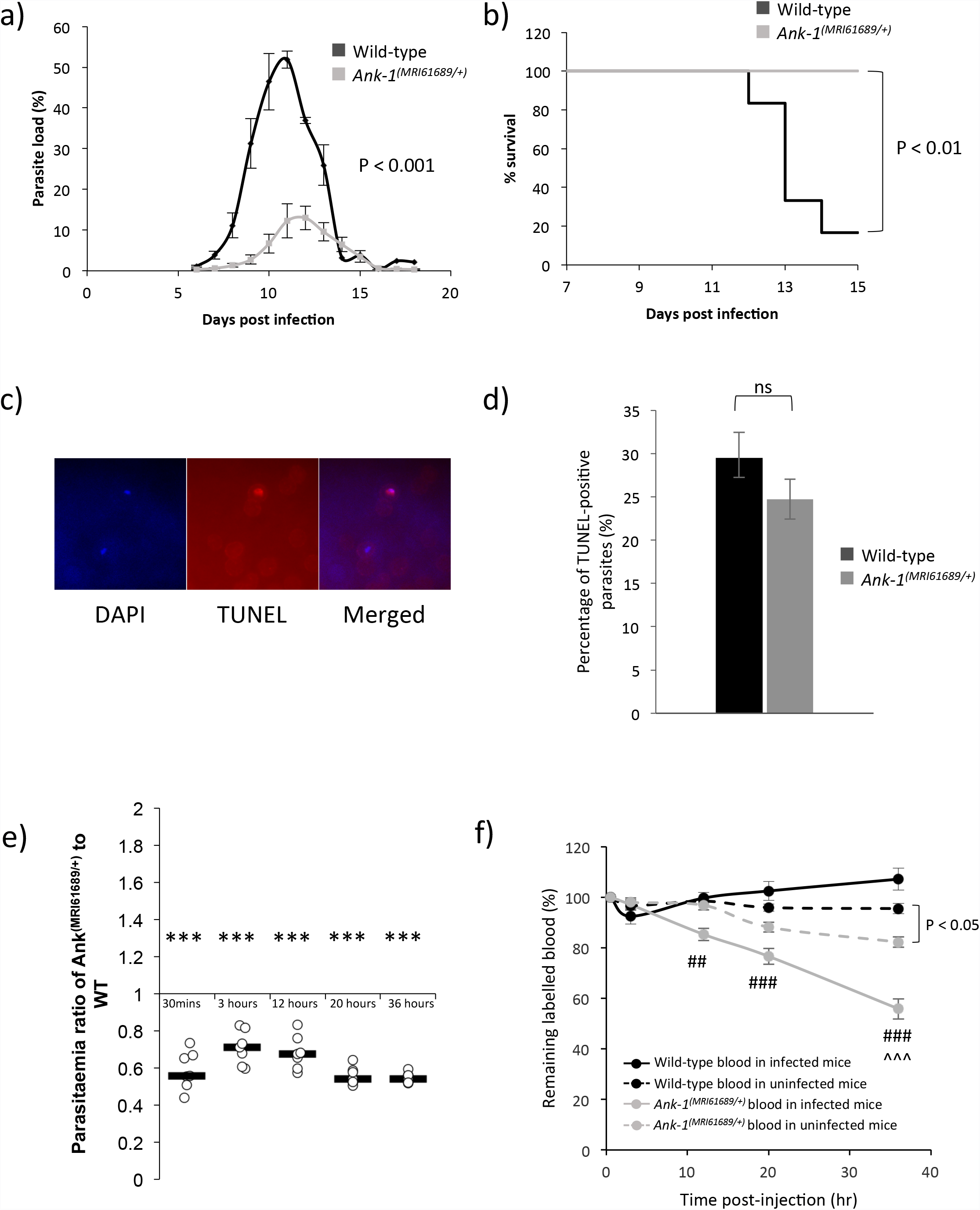
The response of *Ank-1*^*(MRI61689/+)*^ mice to malaria infection. The parasite load of wild-type and *Ank-1*^*(MRI61689/+)*^ mice when infected with 1x10^4^ *P. chabaudi* (a) and the associated survival curve (b). To determine the mechanisms of resistance, parasite intra-erythrocytic growth was assessed through TUNEL assay at 1-10% parasitemia during late trophozoite stage, as visualised from immunofluorescent images showing presence of parasites and TUNEL-positive parasites (c). The number of TUNEL-positive parasites in both wild-type and *Ank-1*^*(MRI61689/+)*^ mice were counted and expressed as a percentage (d) (n=3). For parasite invasion and RBC clearance, The IVET assay was done showing the ratio of infected *Ank-1*^*(MRI61689/+)*^ to wild-type erythrocytes over 36 hours (e) and the relative number *Ank-1*^*(MRI61689/+)*^ and wild-type erythrocyte in both infected and uninfected mice (f) during *P. chabaudi* infection (n=7). ** indicates P<0.01, *** indicates P<0.001, ## and ### indicates P<0.01 and P<0.001 respectively when compared to wild-type RBC number in infected mice, whereas ^^ indicates P<0.01 when compared to *Ank-1*^*(MRI61689/+)*^ RBC number in uninfected mice. Error bars indicate SEM.

### *Ank-1*^*(MRI61689)*^ does not impair the intra-erythrocytic growth of *P. chabaudi*

To elucidate the possible mechanisms of resistance, the effect of the *Ank-1*^*(MRI61689/+)*^ mutation on parasite intra-erythrocytic growth was investigated using the TUNEL assay at 1-10% parasitemia. TUNEL detects DNA fragmentation, a marker for apoptosis or necrosis. In conjunction with a DNA fluorescent dye, DAPI, it is possible to detect dying parasites in the erythrocytes (Figure 4c) ^31,35^. We measured the TUNEL-positivity of *P. chabaudi* in *Ank-1*^*(MRI61689/+)*^ erythrocytes during the late trophozoite stage of the infections, which was the portion of the parasite lifecycle affected by the *Ank1* mutation in the *Ank-1*^*(MRI23420/+)*^ line. As shown in Figure 4d, no differences was observed in the percentage of TUNEL-positive parasites in both wild-type and *Ank-1*^*(MRI61689/+)*^ erythrocytes (24.7 ± 2.3% in *Ank-1*^*(MRI61689/+)*^ mice compared to 29.5 ± 2.9% in wild-type). This indicated that *Ank-1*^*(MRI61689)*^ did not impair parasite intra-erythrocytic growth.

### *Ank-1*^*(MRI61689/+)*^ erythrocyte is resistant to *P. chabaudiinvasion*, and have increased clearance from circulation

Erythrocyte invasion and clearance were assessed via an *in vivo* erythrocyte tracking (IVET) assay. Labelled blood of wild-type and *Ank-1*^*(MRI61689/+)*^ mice were injected into infected wild-type mice during merozoite invasion to examine the ability of *Plasmodium chabaudi* to invade and grow within erythrocytes of both genotypes. The result compared the percentage of parasitized cells of both genotypes and expressed as a ratio of parasitemia in *Ank-1*^*(MRI61689/+)*^ RBC to wild-type RBC populations. As shown in Figure 4e, lower parasitemia ratio (approximately 0.55) was observed from 30 minutes after injection with labelled blood, and was consistently lower in *Ank-1*^*(MRI61689/+)*^ blood over 36 hours post injection, which indicates a lower invasion rate into the *Ank-1*^*(MRI61689/+)*^ RBCs. Additionally, the remaining proportion of labelled RBCs was also monitored over the course of the assay. A significant reduction of *Ank-1*^*(MRI61689/+)*^ erythrocytes in infected mice compared to wild-type erythrocytes was observed, with up to a 45% reduction in *Ank-1*^*(MRI61689/+)*^ RBC number compared with wild-type (Figure 4f). On the other hand, a smaller reduction was observed for *Ank-1*^*(MRI61689/+)*^ erythrocytes compared to wild-type in uninfected mice, suggesting *Ank-1*^*(MRI61689/+)*^ RBCs are more likely to get cleared from circulation during malaria infection. As the percentage of infected erythrocytes was low (5-20-) (Supp. figure 1), this indicates that the majority of the RBCs getting cleared were uninfected RBCs, possibly as a result of bystander effect. This experiment suggested that two possible mechanisms of resistance are both operating to produce the lower parasitemia and increased survival in *Ank-1*^*(MRI61689/+)*^ mice, the reduction of parasite invasion and increased clearance of *Ank-1*^*(MRI61689/+)*^ RBCs.

## Discussion

We report a novel mutation in *Ank-1* gene, MRI61689, causing a hereditary spherocytosis-like phenotype, with reduced MCV, increased osmotic fragility and reduced deformability. MRI61689 is an intronic mutation between exon 17 and 18 where two possible splice variants could arise, one has an introduced acceptor site resulting a frameshift, whereas the other consists of a skipped exon 18. The *Ank-1* mRNA levels were reduced, but no reduction in protein levels were observed. The predicted truncated form (78.5kDa) was also not observed. *Ank-1*^*(MRI61689/+)*^ mice also have increased resistance to *Plasmodium chabaudi* infections, and the erythrocyte invasion was impaired but the intra-erythrocytic growth appeared normal. The *Ank-1*^*(MRI61689/+)*^ RBCs were also more likely to be cleared from circulation during infection, an observation independent of number of parasitized erythrocytes.

In comparison of *Ank-1*^*MRI61689*^ mice to other previously described *Ank-1* mouse models, they appeared comparable to *Ank-1*^*MRI23420*^ mice, but more severe than*Ank-1*^*1674*^ and *Ank-1*^*nb*^ mice. More specifically, most homozygous*Ank-1*^*(MRI61689)*^ mice died within a week after birth, similar to homozygous *Ank-1*^*MRI23420*^ mice, while homozygous *Ank-1*^*1674*^ and *Ank-1*^*nb*^ mice were viable ^30,32^. However, no notable differences were observed in heterozygous *Ank-1*^*MRI61689*^ mice in terms of their RBC microcytosis, morphology and susceptibility to osmotic stress compared to heterozygous*Ank-1*^*1674*^ and *Ank-1*^*MRI23420*^ mice ^31,32^. However, similar to*Ank-1*^*1674/+*^ mice and unlike *Ank-1*^*(MRI23420/+)*^mice,*Ank-1*^*(MRI61689/+)*^ mice exhibited similar levels of ankyrin and other RBC cytoskeletal protein to wild-type, which might suggest compensation by the wild-type allele, and thus warrants further studies.

In humans, many *ANK-1* mutations that result in frameshift have been described, most of which situated in the band 3 binding domain towards the N terminus ^36,37^. While no *Ank-1*^*(MRI61689)*^ homologous mutation has been described in humans, a frameshift mutation has been reported to be in the exon 17, called Ankyrin Osaka I, which gave rise to symptomatic HS ^37^. Furthermore, exon skipping in human *ANK-1* gene has also been documented. Edelman, et al. ^38^ reported a HS patient exhibiting a severe ankyrin-deficient HS due to an introduction of a new splice acceptor site for exon 17, known as ankyrin^*Ankara*^. Under further examination, this mutation was found to give rise to multiple splice forms, including insertions and skipped exons. Most splice forms with insertion were expected to cause frameshift, potentially leading to premature termination of ankyrin ^38^. This finding is in agreement with what we observed for *Ank-1*^*(MRI61689/MRI61689)*^ mice, where frameshift caused by new splice acceptor site, leading to high mortality rate with severe HS-like phenotype in *Ank-1*^*(MRI61689/MRI61689)*^ mice. It is likely that the surviving *Ank-1*^*(MRI61689/MRI61689)*^ mice relied on the exon-skipping splice form to produce in-frame functional *Ank-1* proteins. However, further studies are required for support this hypothesis.

In terms of the response to malaria infection, *Ank-1*^*(MRI61689/+)*^ mice exhibited similar degree of malaria resistance as *Ank-1*^*(MRI23420/+)*^ and *Ank-1*^*1674/+*^ mice, with at least 30-40% reduction in parasitemia and increased survival ^31,32^, and unlike *Ank-1*^*nb/+*^ mice with only 10% reduction ^30^. Previous studies suggested the reduction in erythrocyte invasion and intra-erythrocytic growth to be the major resistance mechanisms ^30,31^. However, normal parasite invasion was reported in *Ank-1*^*1674/+*^ mice ^32^. As a result, we have explored the potential mechanisms of *Ank-1*^*MRI61689/+*^ mice in this study in attempt to elucidate the complex roles ankyrin plays during malaria infections.

First, we report that *Ank-1*^*(MRI61689/+)*^ mice exhibit normal parasite intra-erythrocytic growth (Figure 4d), in contrast to *Ank-1*^*(MRI23420/+)*^ mice. TUNEL assay detects the presence of DNA fragmentation which occurs during apoptosis and necrosis ^39^, which indicates dying parasites in RBCs ^35^. McMorran, et al. ^35^ reported TUNEL-positive parasites in C57BL/6 wild-type mice at a level consistent with the observations in this study. On the other hand, Greth, et al. ^31^ reported a lower TUNEL-positive in their SJL/J wild-type mice, which most likely due to differences in the genetic background of experimental mice. Nevertheless, we did not observe abnormal parasite morphology under light microscopy unlike *Ank-1*^*(MRI23420/+)*^ mice, which support the deductions of parasite death from the TUNEL assays. We also did not observe a difference in gametocyte numbers (Supp. Figure 2), indicating gametocytogenesis was not affected. However, we cannot exclude the possible growth retardation that might occur in other parasite stages, which were not tested in this study.

In terms of parasite invasion, *Ank-1*^*(MRI61689/+)*^ RBCs were found to be more resistant to merozoite invasion as shown in the IVET assay (Figure 4e). The *Ank-1*^*(MRI61689/+)*^ erythrocytes were less infected compared to the wild-type 30 minutes after injection during merozoite invasion, and stayed consistently lower compared to wild-type throughout the erythrocytic cycle. This reduction in invasion has also been observed in *Ank-1*^*(MRI23420/+)*^ mice ^31^. In contrast, Rank, et al. ^32^ reported no difference in parasite invasion for *Ank-1*^*1674/+*^ mice, indicating a different effect mediated by *Ank-1*^*1674*^ mutation compared to *Ank-1*^*(MRI23420)*^ and *Ank-1*^*(MRI61689)*^ mutations.

From these comparisons with other ankyrin mouse models, it is evident that RBC cytoskeleton plays an important yet complex role during malaria infections. However, the exact mechanism for each of these different phenotypes for each ankyrin haplotype remains elusive, different ankyrin mutations can exert different effects on the parasites depending on the location of the mutations, giving rise to multiple resistance mechanisms. This hypothesis is also consistent with the heterogeneous HS symptoms associated with ankyrin mutations, which highlights the complicated interactions between RBC cytoskeletons and malaria parasites.

On the other hand, one important observation from the IVET assay is the rapid clearance of *Ank-1*^*(MRI61689/+)*^ erythrocytes from the circulation within 36 hours post-injection, up to 40% of the initial RBC number (Figure 4f). However, at this timepoint the parasitemia of the host mice was only 5-20% (Supp. Figure 3), therefore the rate of RBC clearance cannot be explained by the clearance of parasitized RBCs, instead, it is likely that the majority of the RBCs being cleared were uninfected. In comparison, no loss was observed for wild-type erythrocytes in both infected and uninfected mice. Since, in these experiments both wild-type and *Ank-1*^*(MRI61689/+)*^ blood were subjected to the same host environment simultaneously, it implies that the clearance of *Ank-1*^*(MRI61689/+)*^ erythrocytes is cell autonomous, rather than due to other effects of the host animal during infectiom. This is further supported by the observation that increased *Ank-1*^*(MRI61689/+)*^ erythrocyte clearance in uninfected mice, indicating that *Ank-1*^*(MRI61689/+)*^ erythrocytes were predisposed for clearance. This is the first observation of increased, uninfected RBC clearance associated with an ankyrin mutation. Bystander clearance is typically observed in inflammation, such as during sepsis ^40^, and is thought to cause severe malaria anaemia during malaria infection in humans through the destruction of normal uninfected RBCs ^41,42^. It is possible that *Ank-1*^*(MRI61689)*^ causes a more exaggerated bystander effect during malaria infection, leading to a further reduction of *Ank-1*^*(MRI61689/+)*^ erythrocyte numbers.

Reduced RBC deformability was proposed to be one of the mechanisms of bystander clearance during malaria anaemia through phagocytosis and splenic filtration ^43,44^. From the *in vitro* spleen retention assay (Figure 1d), *Ank-1*^*(MRI61689/+)*^ erythrocytes exhibited reduced deformability, making them more likely to be retained in the filter layer, which is likely to promote their destruction in *in vivo* settings. Therefore, we proposed that *Ank-1*^*MRI61689*^ mutation causes alteration to the erythrocyte, which renders them more likely to be cleared from the circulation.

In summary, we report that ENU-induced *Ank-1*^*MRI61689*^ mutation causes an HS-like phenotype in mice, and confers significant resistance to *P. chabaudi* infection. We propose that *Ank-1*^*(MRI61689/+)*^ erythrocytes are significantly resistant to parasite invasion but appeared to support normal trophozoite development, although it is possible that this mutation might affect growth of other parasite stages. We also described a novel observation of increased RBC clearance associated with this ankyrin mutation. This study emphasizes the importance of RBC cytoskeletal proteins in mediating multiple complex mechanisms of resistance towards malaria, which provide further insights to the complex interaction between the host and parasites.

## Methods and Materials

### Mice and Ethics Statement

All mice used in this study were housed with 12 hour light-dark cycle under constant temperature at 21 °C. All procedures were conformed to the National Health and Medical Research Council (NHMRC) Australian code of practice. Experiments were performed under ethics agreement AEEC A2014/54, which were approved by the Australian National University animal ethics committees.

### ENU Mutagenesis and Dominant Phenotype Screening

B6.BKS(D)-Lepr^db^/J male mice were injected intraperitoneally with two dose of 100 mg/kg ENU (Sigma-Aldrich, St Louis, MO) at one week interval for a recessive suppressor screen of db/db mutation. The treated males (G0) were crossed to females from the isogenic background to produce the first generation progeny (G1). The seven-week-old G1 progeny were bled and analysed on an Advia 120 Automated Haematology Analyser (Siemens, Berlin, Germany) to identify abnormal red blood cell count. Mouse carrying MRI61689 mutation was identified with a “mean corpuscular volume” (MCV) value three standard deviations lower from other G1 progeny. It was crossed with B6.BKS(D)-Lepr^db^/J mice to produce G2 progeny to test the heritability of the mutations and the dominance mode of inheritance. Mice that exhibited low MCV (<48fL) were selected for whole exome sequencing and genotyping. This abnormal red blood cell count was unrelated to the obesity phenotype.

### Microscopy

For light microscopy, cells were briefly fixed in methanol for one minute and air-dried before being stained in a 10% Giemsa solution (Sigma-Aldrich, St Louis, MO) at pH 7.4 for 10 minutes. For scanning electron microscopy (SEM), fresh blood was first fixed immediately upon drawing in 3% EM-grade glutaraldehyde (Sigma-Aldrich, St Louis, MO) overnight at 4°C. The samples were washed with mouse tonicity phosphate buffered saline (MT-PBS) (150mM NaCl, 16mM Na_2_HPO_4_, 4mM NaH_2_PO_4_, pH 7.4) 3 times, 10 minutes soak each. The cells were then adhered to the cover slips with 0.1% polyethyleneimine (PEI) for 10 minutes, before washing off with MT-PBS. The cells were then dried serially using 30%, 50%, 70%, 80%, 90%, 100%, 100% ethanol, each with 10 minutes soak. The cells were then soaked in 1:1 ethanol: hexamethyldisilazane solution for 10 minutes, followed by 2 washes with 100% hexamethyldisilazane, each 10 minutes. The coverslips were then air-dried overnight and coated with gold and examined under JEOL JSM-6480LV scanning electron microscope.

### Osmotic Fragility Measurement

To assess the susceptibility of RBC membrane to osmotic stress, 5µl of mouse whole blood was diluted 100-fold with phosphate buffer (pH 7.4) containing 0 to 10g/L of sodium, and incubated for at least 10 minutes at room temperature. The cells were centrifuged at 800g for 3 minutes, and the supernatant, which contains free haemoglobin, was measured at 540nm to assess the degree of haemolysis. The absorbance values were expressed as percentage of haemolysis, with haemolysis at 0g/L sodium considered as 100% lysis.

### *In vitro* spleen retention assay

The RBC deformability of both wild-type and *Ank-1*^*(MRI61689/+)*^ were assessed according to the protocol described previous by Deplaine, et al. ^45^ with modifications. Briefly, RBCs from wild-type and *Ank-1*^*(MRI61689/+)*^ mice were stained with 10µg/ml of either hydroxysulfosuccinimide Atto 633 (Atto 633) or hydroxysulfosuccinimide Atto 565 (Atto 565) (Sigma-Aldrich, St Louis, MO), followed by three washes with in MTRC (154mM NaCl, 5.6mM KCl, 1mM MgCl_2_, 2.2mM CaCl_2_, 20mM HEPES, 10mM glucose, 4mM EDTA, 0.5% BSA, pH 7.4, filter sterilized). The stained RBCs were mixed in equal proportion and diluted with unstained wild-type RBCs to give approximately 10-20% of the sample being labelled RBCs. The samples were further diluted to 1-2% haematocrit with MTRC, before passing through spleen retention filter bed. The pre-filtered and post-filtered samples were analysed on BD LSRFortessa (BD Biosciences, Franklin Lakes, NJ) flow cytometer to determine the proportion being retained in the filter bed.

### Whole exome sequencing

DNA from two G2 mice carrying the abnormal red blood cell parameters (MCV <48fL) were extracted with Qiagen DNeasy blood and tissue kit (Qiagen, Venlo, Netherlands) for exome sequencing as previous described ^46^. Briefly, at least 10µg of DNA was prepared for exome enrichment with Agilent Sure select kit paired-end genomic library from Illumina (San Diego, CA), followed by high throughput sequencing using a HiSeq 2000 platform. The bioinformatics analysis was conducted according to the variant filtering method previously described by Bauer, et al. ^47^. Private variants that were shared between the two mutants but not with other B6.BKS(D)-Lepr^db^/J, C57BL/6 mice or previously described ENU mutants were annotated using ANNOVAR ^48^. Private non-synonymous exonic and intronic variants within 20 bp from the exon spicing sites were retained as potential candidate ENU mutations.

### PCR and Sanger sequencing

DNA from mutant mice were amplified through PCR with 35 cycles of 30 seconds of 95^°^C denaturation, 30 seconds of 56-58^°^C annealing and 72^°^C elongation for 40 seconds. The primers used in the PCR are described as below. The PCR products were examined with agarose gel electrophoresis before being sent to the Australian Genome Research Facility (AGRF) in Melbourne, Australia, for Sanger sequencing. Logarithm of odds (LOD) score was calculated based on the number of mice that segregated with the candidate mutations.

### Primers

*Acp5*-F: 5’-CAGAAGGATGCCTTTGGGTA-3’; *Acp5*-R: 5’-ACCAGCGCTTGGAGATCTTA-3’

*Kcnk1*-F: 5’-GGGCCTTTTCCTCCTTACAGA-3’; *Kcnk1*-R: 5’-CAGGAAACGGTGACAAATCC-3’

*Epas1*-F: 5’-GGAAGCCAGAACTTCGATGA-3’; *Epas1*-R: 5’-GTAGTGTTCCCTGGGGTGT-3’

*Picalm*-F: 5’-TCACTGAATGTAATTGGGATATCAT-3’; *Picalm*-R: 5’-CACCCTCTCTTCACTTTTGTG-3’

*Socs6*-F: 5’-CCGCTTTGTTATCCGTCAGT-3’; *Socs6*-R: 5’-TGGCAGCAAAGACTTCAATG-3’

*Ank1*-F: 5’-TCCCTGGCTTAAAGTTGGTG-3’; *Ank1*-R: 5’-CTCTCCCTTAGCTGCATTCC-3’

### Quantitative PCR and cDNA sequencing

RNA was isolated from embryonic livers of E14 embryos using Qiagen RNeasy kit (Qiagen, Venlo, Netherlands), followed by cDNA synthesis using Transcriptor High Fidelity cDNA Synthesis Kit (Roche, Basel, Switzerland). Quantitative PCR was carried out on ViiA™ 7 Real-Time PCR System (Thermo Scientific, Waltham, MA). The ΔΔC_T_ method ^49^ was used to determine the cDNA levels of *Ank-1* and the housekeeping gene β-actin and expressed as a fold-change of the mutants to the wild-type. The primers used for *Ank-1* gene spanned exon 2 to 4: *Ank-1*-F: 5’-TAACCAGAACGGGTTGAACG-3’; *Ank-1*-R: 5’-TGTTCCCCTTCTTGGTTGTC-3’; β-Actin-F: 5’- TTCTTTGCAGCTCCTTCGTTGCCG-3’; β-Actin-R: 5’- TGGATGCGTACGTACATGGCTGGG-3’.

To characterize the effect of MRI61689 mutation, cDNA were amplified through PCR using two primers set were design as shown below: Primer set 1 was designed to amplify wild-type *Ank-1* transcript, whereas primer set 2 was designed to only amplify the predicted mutant transcript with 11bp insertion. Amplified PCR product were analysed using agarose gel electrophoresis and each band was purified and sequenced.

Primer set 1: Forward: ATGCAGAGTCGGTACAAGGC; Reverse: CCGTTCGAGCTGACCTCATT

Primer set 2: Forward: CCTGGGGAACAAGTTTCTTT; Reverse: GTGCAAGGGGCTGTATCCTA

### SDS-PAGE, Coomassie staining and Western blot

RBC ghosts were prepared by lysing mouse RBCs with ice-cold 5mM phosphate buffer (ph7.4) and centrifuged at 20,000g for 20 minutes followed by removal of the supernatant. The pellet was further washed with the 5mM phosphate buffer until the supernatant became clear. The RBC ghosts or whole blood lysates were denatured in SDS-PAGE loading buffer (0.0625M Tris pH 6.8, 2% SDS, 10% glycerol, 0.1M DTT, 0.01% bromophenol blue) at 95°C for 5 minutes before loading onto a Mini-PROTEAN^®^ TGX^TM^ Precast Gels (Bio-Rad, Hercules, CA). The gels were then either stained with Coomassie blue solution (45% v/v methanol, 7% v/v acetic acid, 0.25% w/v Brilliant Blue G) overnight or transferred to a nitrocellulose membrane. The western blot was carried out using these primary antibodies: anti-alpha 1 spectrin (clone 17C7), anti-beta 1 spectrin (clone 4C3) (Abcam, Cambridge, UK), anti-GAPDH (clone 6C5) (Merck Millipore, Darmstadt, Germany), anti-N-terminal *Ank-1* “p89”, anti-Band 3 and anti-protein 4.2 (kind gifts from Connie Birkenmeier, Jackson Laboratory, US).

### Malaria infection

250µl of thawed *P. chabaudi adami* infected blood was injected into the intraperitoneal cavity of a C57BL/6 donor mouse. When the donor mouse reached 1-10% parasite load (parasitemia), blood was collected through cardiac puncture. The parasitized blood was diluted with Krebs’ buffered saline with 0.2% glucose as described previously ^50^. Each experimental mouse was infected with 1x10^4^ parasites intraperitoneally. The parasitemia of these mice were monitored either using light microscopy or flow cytometry.

### Terminal deoxynucleotidyl transferase dUTP nick end labelling (TUNEL) staining

3µl of infected blood containing 1-10% parasitemia were collected during trophozoite stage and fixed in 1 in 4 diluted BD Cytofix^TM^ Fixation Buffer (BD Biosciences, Franklin Lakes, NJ) for at least day until they were needed. Each sample was then washed twice with MT-PBS, and adhered to a glass slide pre-coated with 0.1% polyethylenimine (PEI) for 10 minutes at room temperature. The excess cells were washed off with the wash solution from APO-BrdU TUNEL assay kit (Thermo Scientific, Waltham, MA) and incubated overnight at room temperature with TUNEL labelling solution (1mM Cobalt Chloride, 25mM Tris-HCl pH 6.6, 200mM sodium cacodylate, 0.25mg/ml BSA, 60uM BrdUTP, 15U Terminal transferase). The slides were washed three times with rinse buffer from APO-BrdU TUNEL assay kit, followed by staining with 50µg/ml of anti-BrdU-Biotin antibody (Novus Biologicals, Littleton, CO) in MT-PBT (MT-PBS, 0.5% BSA, 0.05% Triton X-100) for 1 hour. The slides were then washed three times with MT-PBT, followed by probing with 2µg/ml Alexa Fluor^®^ 594 conjugated streptavidin (Thermo Scientific, Waltham, MA). Next, they were washed three times with MT-PBS and mounted with SlowFade^®^ Gold antifade reagent with DAPI (Thermo Scientific, Waltham, MA) and sealed. When the slides were dried, they were examined using Axioplan 2 fluorescence light microscope (Carl Zeiss, Oberkochen, Germany) between 600x to 1000x magnification. At least 100 DAPI-positive cells were counted, and each was graded as either positive or negative for TUNEL staining, as an indication of DNA fragmentation.

### *In vivo* erythrocyte tracking (IVET) assays

The IVET assay was carried out as previously described by Lelliott, et al. ^51,52^. Briefly, 1.5ml to 2ml of whole blood was collected from wild-type and *Ank-1*^*(MRI61689/+)*^ mice via cardiac puncture. Both wild-type and *Ank-1*^*(MRI61689/+)*^ blood were either stained with 10µg/ml of Atto 633 or 125µg/ml of EZ-Link™ Sulfo-NHS-LC-Biotin (Biotin) (Thermo Scientific, Waltham, MA) for 45 minutes at room temperature, followed by washing three times with MT-PBS. The *Ank-1*^*(MRI61689/+)*^ blood was mixed with wild-type blood in two different dye combinations to correct for any dye effects. 1x10^9^ erythrocytes were injected intravenously into infected wild-type mice at 1-5% parasitemia during schizogony stage, usually 8-10 days post-infection with 1x10^4^ parasites. Blood samples were collected at 30 minutes, 3hours, 12 hours, 20 hours and 36 hours after injection. The ratio of infected *Ank-1*^*(MRI61689/+)*^ to wild-type erythrocytes was determined on flow cytometry, as an indication of the relative susceptibility to malaria infections between wild-type and *Ank-1*^*(MRI61689/+)*^ mice. The proportion of labelled blood populations were also tracked over time to determine the clearance of these RBCs from the circulation.

### Flow cytometry analysis of blood samples

For both malaria infections and IVET assay, 2µl of whole blood samples were stained with 2μg/ml streptavidin-PE-Cy7 (only for experiments with biotinylated erythrocytes), 1μg/ml anti-CD45– allophycocyanin (APC)–eFluor 780 (clone 30-F11), 1μg/ml anti-CD71 (TFR1)–PerCP–eFluor 710 (clone R17217) (eBioscience, San Diego, CA), 4µM Hoechst 33342 (Sigma-Aldrich, St Louis, MO) and 12µM JC-1 (Thermo Scientific, Waltham, MA) in MTRC. All samples analysed through flow cytometry were performed on BD LSRFortessa (BD Biosciences, Franklin Lakes, NJ), where 100,000 to 2,000,000 events were collected and visualized on FACSDiva^TM^ and FlowJo software. The RBCs and leukocytes were first selected on forward scatter and side scatter channels (FSC/SSC) signals, followed by gating of single cells based on FS area to height ratio. RBCs were further isolated by gating on CD71 negative and CD45 negative population, followed by gating on Atto-labelled and Biotin-labelled erythrocytes on appropriate channels (APC for Atto-633, PE for Atto-565 and PE-Cy7 for Biotin). The parasitemia of each labelled erythrocyte population was determined by gating on Hoechst 33342 positive and JC-1 positive population.

### Statistical analysis

The LOD score method coupled with Bonferroni correction was used to determine the causative mutation for MRI61689. The statistical significance of the malaria survival was tested using the Log-Rank test. The statistical significance of parasite infection was determined via the statmod software package for R (http://bioinf.wehi.edu.au/software/compareCurves) using the ‘compareGrowthCurves’ function with 10,000 permutation, followed by adjustments for multiple testing. The statistical significance for the ratios of IVET assays was determined using the one sample t-test with hypothetical mean of 1. For the rest of the results, statistical significance was determined using two-tailed Students t-tests.

## Acknowledgement

We would like to acknowledge Shelley Lampkin and Australian Phenomics Facility (APF) for the maintenance of the mouse colonies. This study was funded by the National Health and Medical Research Council of Australia (Program Grant 490037, and Project Grants 605524 and APP1047090), Australian Society of Parasitology (ASP), OzEMalaR, National Collaborative Research Infrastructure Strategy (NCRIS), the Education Investment Fund from the Department of Education and Training, the Australian Phenomics Network, Howard Hughes Medical Institute and the Bill and Melinda Gates Foundation. We would finally like to thanks two anonymous reviewers for their insightful comments on this manuscript.

---

**Supplementary figure 1.**
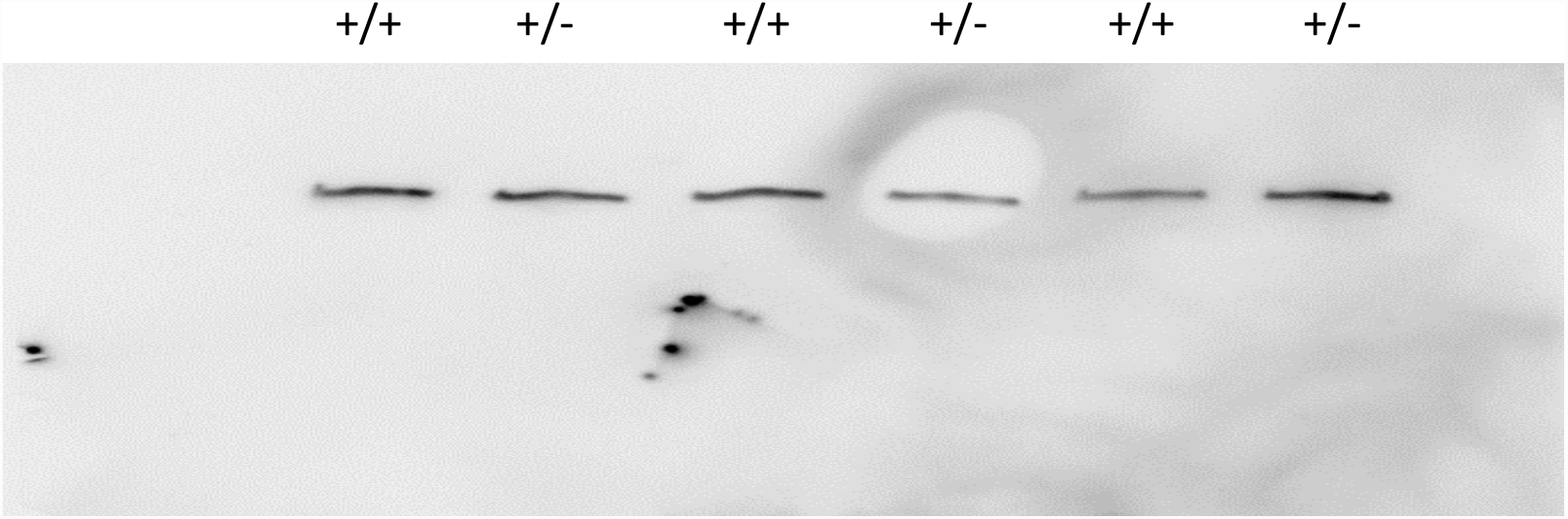
The full western blot membrane when probed with anti-beta-spectrin antibody. The disparity of band intensity as shown in Figure 3c is due to uneven ce of the membrane rather than post6processing issue.

**Supplementary figure 2.**
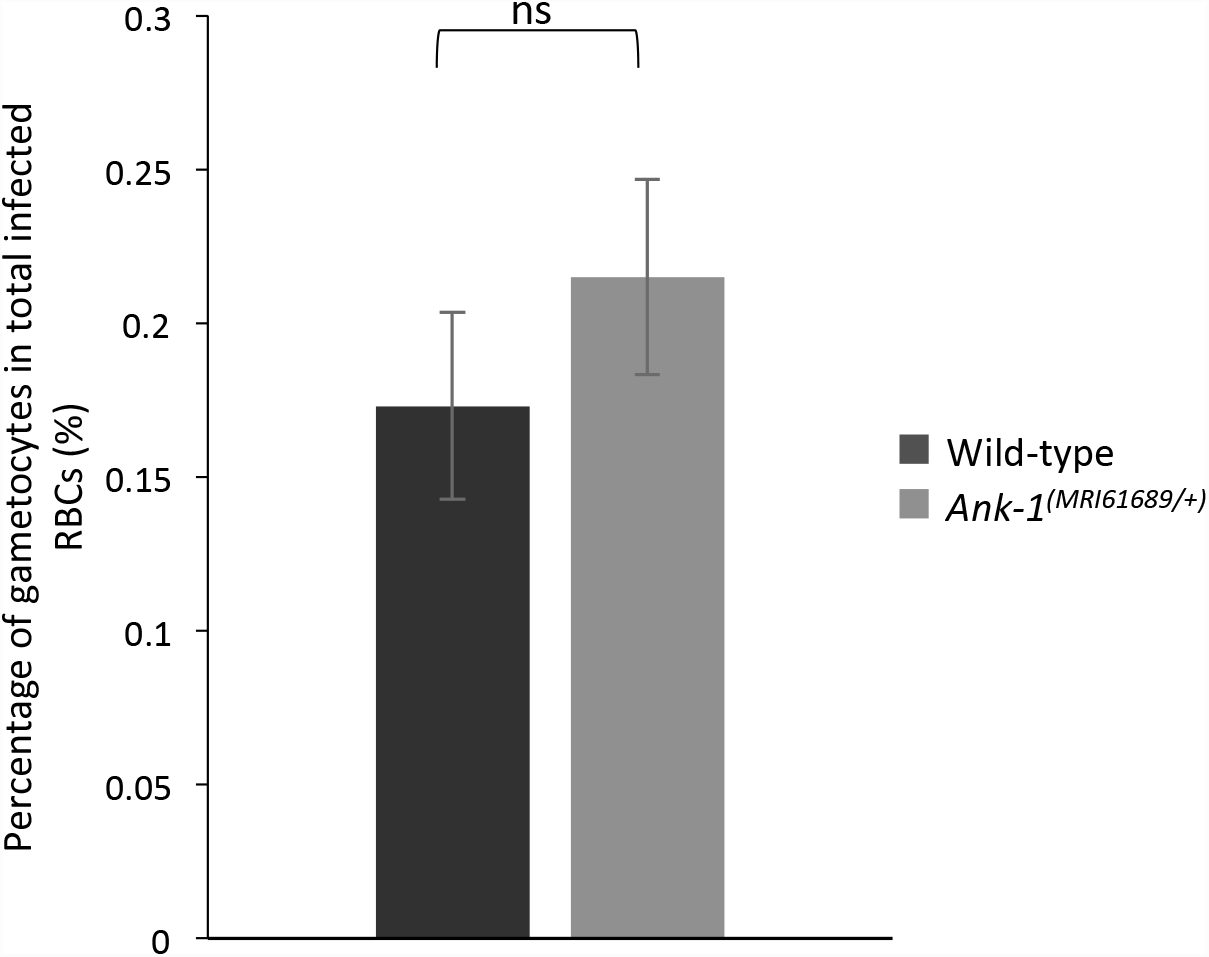
The percentage of gametocytes of wild-type and of *Ank-1*^*(MRI61689/+)*^ during malaria infection. Parasite gametocyte numbers were counted under light microscopy at 15–30% parasitaemia, and the proporUon of gametocytes to total infected RBCs calculated (n=6). Error bars indicate SEM.

**Supplementary figure 3.**
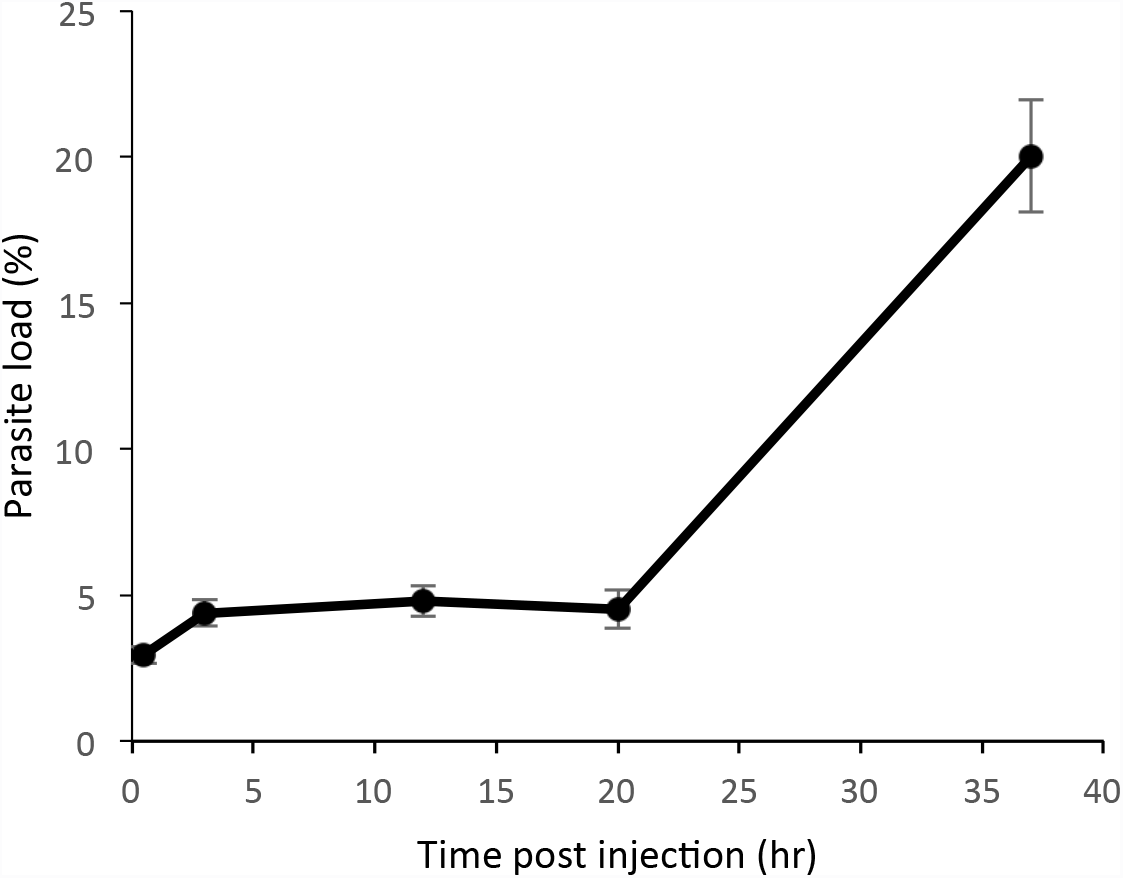
The parasite load of the mice during IVET assays. The parasite load host mice during IVET assay (n=7). Error bars indicate SEM.

